# Understanding requires tracking: noise and knowledge interact in bilingual comprehension

**DOI:** 10.1101/609628

**Authors:** Esti Blanco-Elorrieta, Nai Ding, Liina Pylkkänen, David Poeppel

**Affiliations:** Department of Psychology, New York University, New York, NY, USA; NYUAD Institute, New York University Abu Dhabi, Abu Dhabi, UAE; Speech, Cognition and Neuroscience Department, Zhejiang University, Zhejiang Sheng, China; Department of Linguistics, New York University, New York, NY, USA; Department of Neuroscience, Max Planck Institute for Empirical Aesthetics, Frankfurt, Germany; Center for Neural Science, New York University, New York, NY, USA; Center for Language, Music, and Emotion, New York University, New York, NY, USA

## Abstract

Understanding speech in noise is a fundamental challenge for speech comprehension. This perceptual demand is amplified in a second language: it is a common experience in bars, train stations, and other noisy environments that degraded signal quality severely compromises second language comprehension. Through a novel design, paired with a carefully selected participant profile, we independently assessed signal-driven and knowledge-driven contributions to the brain bases of first versus second language processing. We were able to dissociate the neural processes driven by the speech signal from the processes that come from speakers’ knowledge of their first versus second languages. The neurophysiological data show that in combination with impaired access to top-down linguistic information in the second language, the locus of bilinguals’ difficulty in understanding second language speech in noisy conditions arises from a failure to successfully perform a basic, low-level process: cortical entrainment to speech signals above the syllabic level.

**Significance statement:** Over half of the world’s population is multilingual. Although proficiency in a second language improves over time, even to the point of reaching native-like proficiency, one persistent difficulty is understanding a second language in noisy environments (e.g. in crowds, train stations, etc.). We examined this processing impairment using a novel frequency-tagging paradigm with magnetoencephalography (MEG). We found that the reason underlying bilinguals’ difficulty in understanding second language speech in noisy conditions is induced by a failure to neurally track the operations reflecting structure (i.e., phrase) building.

## Introduction

Speaking more than one language is the norm for the majority of the world’s population (Craik & Bialystok, 2006; US Census Bureau, 2010), and multilingualism is increasing notably (Cenoz et al., 2006). Although language proficiency can improve remarkably through exposure over the years, even to the point of reaching native-like proficiency, there is a familiar phenomenon that remains challenging throughout the life of a bilingual individual: in noisy environments, comprehension is hard in a second language - but seems relatively effortless in a first. Our understanding of the computational and neural foundations of this ubiquitous phenomenon is rather limited. A few hypotheses have attempted to account for this experience (Ferreira, Engelhardt, & Jones, 2009; Golestani, Hervais-Adelman, Obleser, & Scott, 2013; Hahne & Friederici, 2001; Hervais-Adelman, Pefkou, & Golestani, 2014), and although somewhat different in scope, they have all proposed a lack of successful use of top-down linguistic information as the source of this effect. The rationale is as follows: in any given situation where humans listen to a degraded signal, they use top-down linguistic knowledge such as the sentential context or predicted semantic meaning of the sentence to calculate and repair the message that has been obscured by the poor-quality of the input. Researchers have argued that bilingual individuals not having as easy an access to this top-down semantic information in their second language leads to an inability to repair the speech signal and to consequently not understand the message (Hervais-Adelman et al., 2014). Here we tested the possibility that in addition to a failure to accurately apply high-level linguistic information, the source of this persistent difficulty may also lie in an inability to perform a lower level process reported to aid comprehension (Zoefel, Archer-Boyd, & Davis, 2018): the neurophysiologically well-established concept of neural entrainment to speech (Buzsáki, & Draguhn, 2004; Lakatos, Karmos, Mehta, Ulbert, & Schroeder, 2008).

In order to characterize quantitatively the effect of noise across different L2 proficiency levels, we recruited bilingual (Mandarin Chinese, American English) participants who were (i) Mandarin Chinese native speakers with low English proficiency, (ii) Mandarin native speakers with high English proficiency (these participants lived in China until adulthood and had learned English since they were young, but only in an academic setting), and (iii) native speakers of Mandarin who were English dominant (born to at least one Mandarin speaking caregiver in an English speaking country). Thus, our carefully selected participant sample covered the full spectrum of possible language proficiency combinations in both languages. We recorded electrophysiological (MEG) responses while participants listened to 4 word sentences at different signal-to-noise (SNR) ratios, varying from completely clear to fully unintelligible speech. We discovered that the neural responses that track the physical speech rhythm are affected by noise – but not by language proficiency. In contrast, responses tracking linguistic structure reflect the interaction between noise and knowledge. Hence, complementing previous research suggesting that greater availability of top-down linguistic information may account for the difference in L1 versus L2 comprehension (Hervais-Adelman et al., 2014), the data show that an automatic lower-level mechanism tracking speech also contributes to the prevalent effect of impoverished comprehension of L2 speech in noise.

## Materials and methods

### Participants

Fifty-one right-handed Mandarin-English bilingual individuals participated in the experiment (16 male, 35 female, mean 20; SD ± 2.45 years). In order to meaningfully characterize the effect of noise across varied second language proficiency levels, we selected Mandarin-English bilinguals with diverse language backgrounds. 16 of the participants were native speakers of Mandarin who had acquired English later in life and had always lived in a Mandarin-speaking environment (age of acquisition Mandarin = 1.31 (1.53 SD), English = 7.62 (3.5 SD)). Their self-reported oral (speaking and understanding) proficiency was 96.8% in Mandarin (4.7 SD) and 68.5% in English (7.1 SD), and their score in the Woodcock-Muñoz English language survey was 51.8%. English had never been their language for socializing, and they had rarely used it outside of a classroom context. 17 participants were native speakers of Mandarin with high proficiency in English. They had acquired English earlier in life at international schools but had grown up in a Mandarin-speaking environment (age of acquisition Mandarin = 1.33 (0.88 SD), English = 5.91 (4.42 SD)). They had moved to the US for undergraduate education at some point in the past 3 years and had since been in an English-speaking environment. Their self-reported oral proficiency was 94.1% in Mandarin (3.2 SD) and 82.9% in English (6.4 SD), and their score in the Woodcock-Muñoz English language survey was 68.8%. Lastly, we tested a group of English dominant speakers (n=18), who were born to at least one Mandarin speaking parent in the US (age of acquisition Mandarin = 1.83 (0.70 SD), English = 2.25 (4.24 SD)). Hence, they had learned Mandarin from birth but the dominant language in their environment and everyday use had always been English. Their self-reported oral proficiency was 76.4% in Mandarin (6.1 SD) and 95.6% in English (2.41 SD), and their score in the English language survey was 91.4%. In this last group, participants reported that their life unfolded fully in English except for at home, where they spoke Mandarin, and they reported their English to be significantly better than their Chinese.

Our grouping criterion was validated post-hoc by submitting participants to K-means clustering based on all the collected language profile variables (i.e., age of acquisition, exposure, self-reported proficiency and quantitative measures of proficiency) and showing that our criterion matched the output of this unsupervised clustering algorithm (t(49) = 7.22; *p* < .001; see Additional materials for detailed language background information). Information about their language use and proficiency level was gathered with a modified version of the language background questionnaire of Marian and colleagues (2007; see additional materials for full language background information). All subjects were neurologically intact with normal or corrected-to-normal vision and all provided informed written consent following NYU Institutional Review Board protocols.

### Stimuli

Participants listened to four-syllable sentences, concatenated and isochronously presented, in English and Mandarin. Mandarin stimuli were 50 four-syllable sentences taken from Ding, Melloni, Zhang, Tian, and Poeppel (2016; MandarinMaterials, Supplementary Table 1, *Four-syllable sentences*), in which the first two syllables formed a noun phrase and the last two formed a verb phrase (Fig. 1B left panel). The combination of the noun and the verb phrase formed a complete sentence. All syllables were presented isochronously, lasted between 75 and 354 ms (mean duration 224 ms; Fig. 1B, top right panel), and were adjusted to 250 ms by truncation or padding silence at the end. Each trial consisted of the sequential presentation of 10 of these sentences, and, crucially, no acoustic gaps were inserted between sentences, as such a gap would constitute an unwanted acoustic cue for segmentation. For this reason, the intensity of the stimulus as shown by the sound envelope, only fluctuated at the syllabic rate (Fig. 1AB, bottom right panels, Supplementary Figure 1 for stimulus spectrograms). That being said, the sequences of four syllables constituted clearly segmentable sentences. English materials consisted of 60 four-syllable sentences. Each sentence consisted of 4 monosyllabic words combined to form two-word noun phrases (adjective + noun) and two-word verb phrases (verb + noun). The combination of these two phrases resulted in four-word sentences (e.g., “big rocks block roads”; Fig. 1A, left panels). All syllables were between 250 and 347 ms in duration and were adjusted to 320 ms by padding or truncation (Fig. 1A top right panel). For both English and Mandarin, a 25-ms cosine window smoothed the offset of each syllable. All sentences in English and Mandarin are displayed in Supplementary materials. Even though the syllable duration in the two languages was different, the manipulation was designed to ensure isochronous presentation, which is the essential feature of the experimental design.

**Fig. 1.**
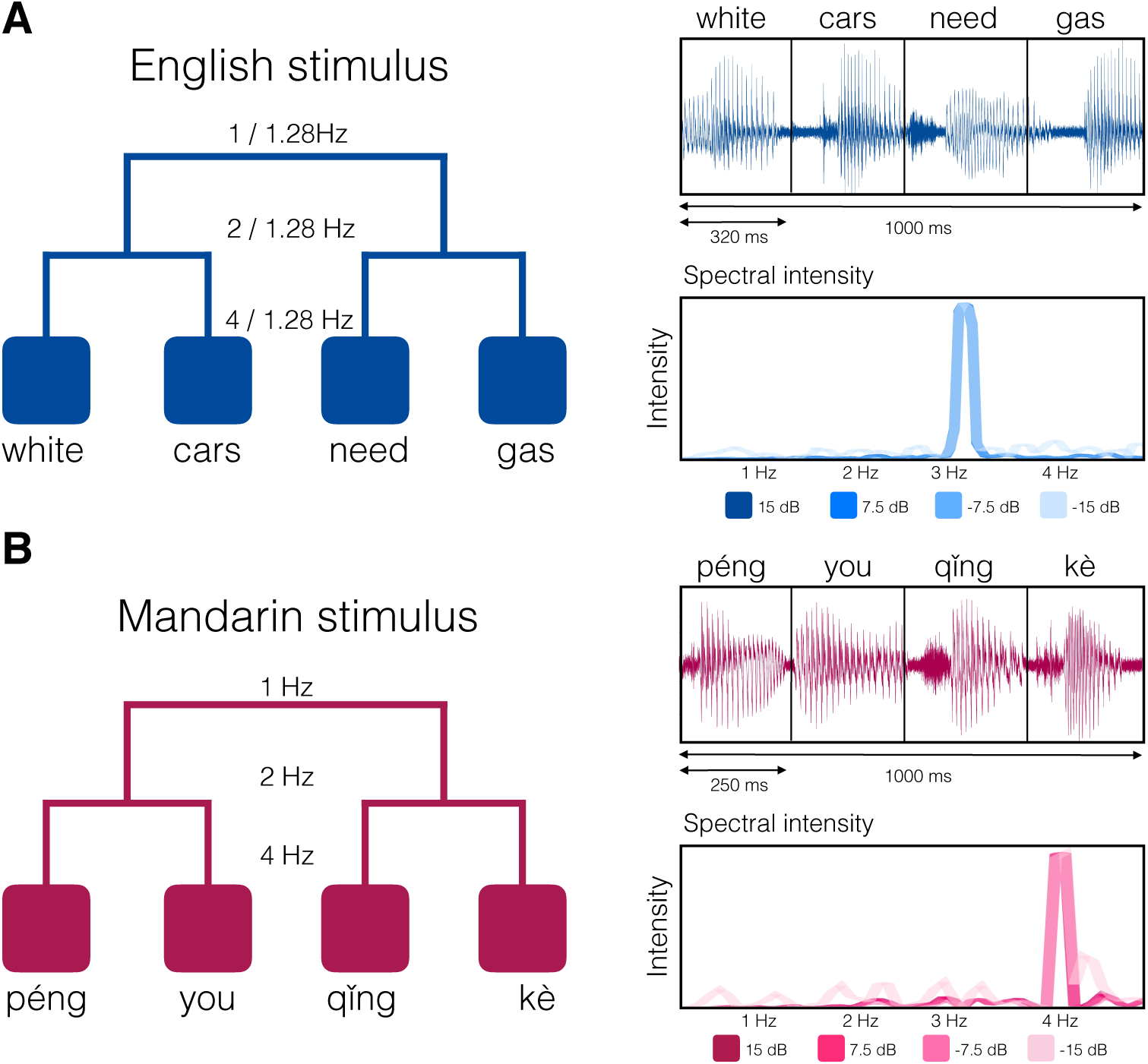
Sample English (A) and Mandarin (B) stimuli. Monosyllabic words were presented isochronously, forming phrases and sentences. Left panels: presentation rate and syntactic structure of the stimuli. In each of A and B, top right: waveform of a sample stimulus. Bottom right: spectral intensity of the stimulus at each tested level of noise, revealing a syllabic-scale rhythm at all levels of noise, but no phrasal-rate or sentential-rate modulation.

We embedded all sentences in 4 different levels of white noise. We first measured the power of the sentences in isolation and then added white noise to reach the desired output signal-to-noise ratio in dB. The SNR levels ranged from +15 dB (clear speech) to −15 dB (unintelligible speech in noise) in 7.5 dB intervals. Although ideally (and eventually) babbling noise may be a better background noise to mirror the type of noise experienced in real life, white noise was selected for a first characterization to avoid confounding semantic, phonological, or language interference effects that went beyond the effects of noise qua noise. Participants heard 160 sentences at each noise level for each language, and the experiment took approximately one hour to complete. Across participants, each four-syllable sentence was presented an equal number of times at each level of noise.

### Procedure

Before the MEG recording, each subject’s head shape was digitized using a Polhemus dual-source handheld FastSCAN laser scanner. MEG data were collected in the Neuroscience of Language Laboratory at NYU using a whole-head 157 channel axial gradiometer system (Kanazawa Institute of Technology, Kanazawa, Japan) as subjects lay in a dimly lit, magnetically shielded room. Trials began with the binaural auditory presentation of the stimuli. Participants listened to sets of 10 randomly presented 4-syllable sentences in either language and were then presented with a 1 – 4 scale onscreen. Listeners had to indicate via button press how much they understood (1 = “nothing at all” and 4 = “everything”). The validity of this comprehension measure has been previously established by showing qualitatively comparable results between this subjective measure of comprehension and participants’ performance on recall tests (Ghitza, 2012; Doelling, Arnal, Ghitza, & Poeppel, 2014). After the button press the next trial began. Following the MEG recording, all participants completed the Woodcock-Muñoz Language Survey to evaluate their proficiency in English. Subjects completed the first four parts of this survey, aimed to assess their oral, listening, reading, and writing skills. The completion of this test took around 45 minutes.

### Data acquisition and preprocessing

MEG data were recorded at 1000 Hz (200 Hz low-pass filter), noise reduced via the continuously adjusted least-squares method (Adachi et al., 2001) in MEG Laboratory software (Yokogawa Electric and Eagle Technology) and epoched from beginning to end of the auditory stimulus. The MEG responses were decomposed into components using a Denoising Source Separation technique (DSS; de Cheveigné & Simon, 2008; for a detailed explanation see Ding et al., 2016), and the first 5 components were kept for analysis and projected back into sensor space. This technique decomposes MEG recordings to extract the neural response components that are consistent over trials, and it was applied to accurately estimate the strength of neural activity phase-locked to the stimulus. To avoid the transient response at the beginning of each trial, data were only analyzed from the beginning of the second sentence of each 10-sentence trial. Single trial responses per noise level and participant were Fourier transformed (DFT) into the frequency domain, and subsequently averaged within condition to obtain an evoked response per condition per participant.

Data were source localized with MNE-Python (Gramfort et al., 2013, 2014). To estimate the distributed electrical current image in the brain at each time sample, we used the Minimum Norm Approach (Hämäläinen and Ilmoniemi, 1994) via MNE (MGH/HMS/MIT Athinoula A. Martinos Center for Biomedical Imaging, Charleston, MA). The cortical surfaces were constructed using an icosahedron subdivision of 4 and mapping an average brain from FreeSurfer (CorTech and MGH/HMS/MIT Athinoula A. Martinos Center for Biomedical Imaging) to the head-shape data gathered from the headscanning process. This generated a source space of 5124 points for each reconstructed surface, leaving ∼6.2 mm of spacing within sources (cortical area per source = ∼39 mm^2^). Then, the boundary-element model method was used to calculate the forward solution. The 100 ms pre-stimulus period was used to construct the noise covariance matrix and to apply as a baseline correction. The inverse solution for each subject was then computed from the noise-covariance matrix, the forward solution, and the source covariance matrix, and was applied to the evoked response for each condition. The application of the inverse solution determined the most likely distribution of neural activity in source space. Minimum norm current estimates were computed for three orthogonal dipoles, of which the root mean square was retained as a measure of activation at that source (thus, the orientation of the dipole was free unsigned). The resulting minimum norm estimates of neural activity were transformed into normalized estimates of noise at each spatial location using the default regularization factor (SNR = 3). Hence, we obtained noise-normalized statistical parametric maps (SPMs), which provide information about the statistical reliability of the estimated signal at each location in the map with millisecond accuracy. Then, those SPMs were converted to dynamic maps (dSPMs). To quantify the spatial resolution of these maps, the point-spread function for different locations on the cortical surface was computed. The point spread is defined as the minimum norm estimate resulting from the signals coming from a current dipole located at a certain point on the cortex. The calculation of the point-spread function following the approach of Dale et al. (2000) reduces the location bias of the estimates, in particular, the tendency of the minimum norm estimates to prefer superficial currents (i.e., their tendency to misattribute focal, deep activations to extended, superficial patterns). Hence, by transforming our minimum norm estimates to dSPM, we obtained an accurate spatial blurring of the true activity patterns in the spatiotemporal maps (Dale et al., 2000).

### Analyses

#### Behavioral data

For the main statistical tests, we conducted mixed-effect model analyses using the lme4 package (Bates, Maechler, & Bolker, 2012) in R (R Core Team, 2012) using Noise (−15 dB, −7.5 dB, 7.5 dB and 15 dB), Proficiency, and the interaction between them as fixed effects and subject as a random effect. Additionally, we conducted complementary categorical analyses within each group of participants to assess the effect of noise in intelligibility within each language profile specifically. For this analysis, responses for each trial were averaged within participant for each noise level. We subsequently conducted a related samples two-tailed t-test across participants to assess whether their comprehension in English and Mandarin at each noise level significantly differed. All reported p-values are FDR corrected for multiple comparisons.

#### Tracking analyses

MEG activity for each trial was averaged within participant at each noise level in the frequency domain. We then subjected the amplitude at each peak to the same linear mixed-effects model used on behavioral data, with Noise, Proficiency and their interaction as fixed effects and subject as a random effect. Having assessed that the interaction of noise and proficiency significantly accounted for the amplitude of the peaks, we split participants by proficiency to unpack the nature of this effect and assess the effect of noise at both the syllabic and phrasal peak within these groups. For each spectral peak, a one-tailed paired t-test was used to test if the neural response across all sensors in a frequency bin was significantly stronger across participants than the average of the four neighboring frequency bins (two bins on each side). We corrected for multiple comparisons across t-tests using FDR-correction. The application of a 1000-permutation test in lieu of the original t-tests revealed the same significant results.

#### Source localization analysis

Having obtained distinct tracking effects across languages for each participant group at −7.5 dB, we turned to the source localized data to identify from where in the cortex this activity was emerging. For this purpose, we subtracted the response to the English sentences at −7.5 dB from the response to Mandarin sentences at −7.5 dB (averaged across participants). This analysis hence revealed the localization of the oscillatory analysis effect.

## Results

### Behavioral results

Behavioral results showed that the influence of noise on the comprehension of speech varied based on individuals’ language proficiency. A linear mixed-effects regression model regressing noise, age of acquisition, and language proficiency on comprehension revealed that (i) in addition to noise and proficiency significantly influencing comprehension independently, (ii) they also interacted significantly, such that the decrease in comprehension due to noise was greater the lower the proficiency of the individual in that language. This effect held both for Chinese (Noise: F(1,4551)=148; *p* < .001); Proficiency: F(1, 46.8) = 31.5; *p* < .001); interaction between Noise and Proficiency F(1,4550) = 4.45; p = .03)) and for English stimuli (Noise: F(1,4975)=44.1; *p* < .001); Proficiency: F(1, 47) = 7.72; *p* = .007); interaction between Noise and Proficiency F(1,4975) = 23.39; *p* < .001)).

Next, to unpack these results, we complemented the continuous regression analysis with categorical analyses wherein we assessed the influence of noise in comprehension for each language group. We found that for Mandarin-dominant participants with low English proficiency (Fig. 2A), the lower comprehension of English compared to Mandarin was constant across all levels of noise (*p* < .001). However, Mandarin speakers with high English proficiency understood both languages equally in clear speech (15 dB *p* = .09; 7.5 dB *p* = .14), but their comprehension of English was severely impaired in noisy conditions (−7.5 dB and −15 dB *p* < .001; Fig. 2B). Finally, we found that English dominant bilinguals showed an overall impaired comprehension of Mandarin, except at the highest level of noise; in that case, participants did not report understanding in either language (15 dB; 7.5 dB, −7.5 dB *p* < .001; −15 dB *p* = .12; Fig. 2C). Overall, behavioral results reveal distinct interactions between noise and comprehension, contingent on language knowledge.

**Fig. 2.**
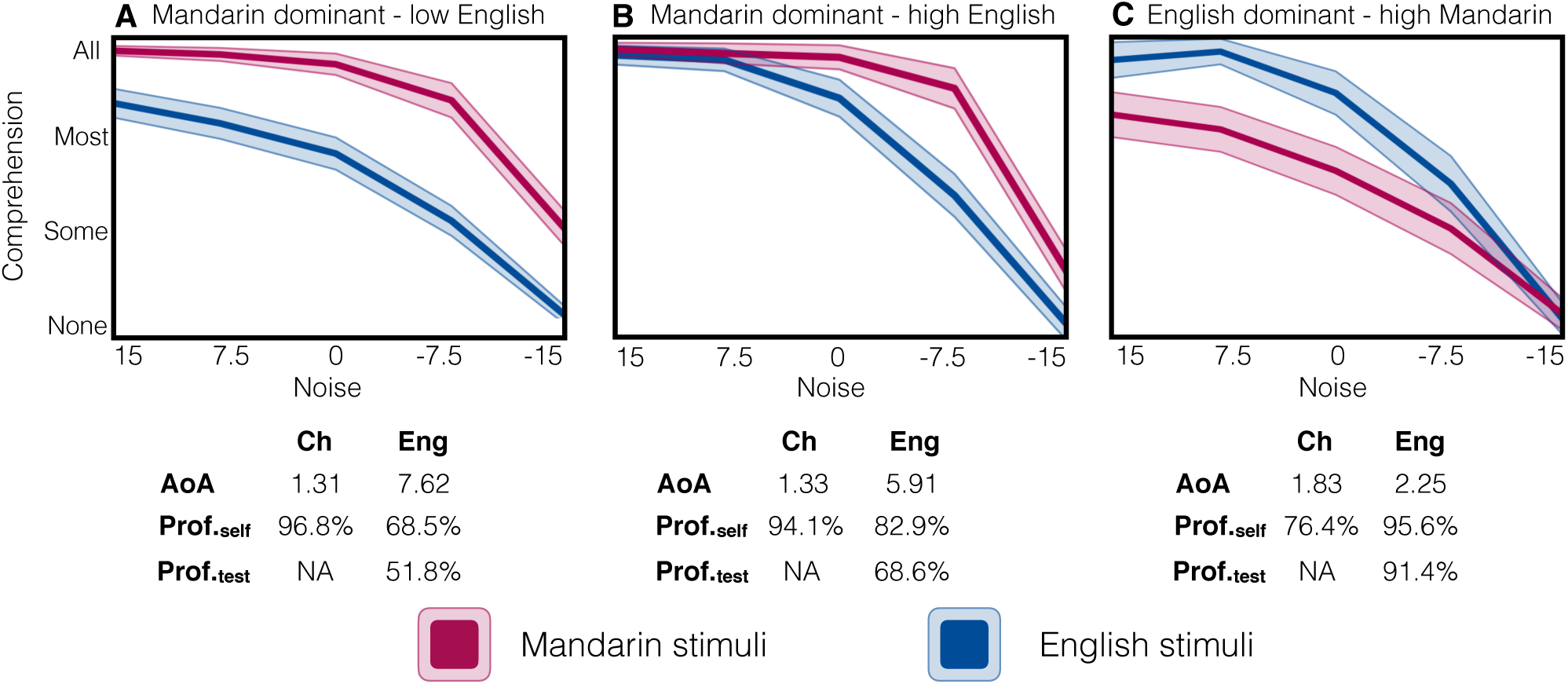
Comprehension performance for participants as a function of noise, averaged across subjects (shaded area: 95% C.I.). Below each graph, we report the average age of acquisition (AoA), self-reported proficiency (Prof._self_) and English proficiency score in the Woodcock language questionnaire (Prof._test_) for the participants in each group.

### MEG: cortical tracking results

While participants listened to the sentences, we recorded neural activity with MEG. Cortical oscillations have been proposed to be likely candidates for segmentation of continuous speech (Gross et al., 2013; Luo & Poepple, 2007; Zoefel et a., 2018). We performed analyses aimed to quantify how entrainment to different linguistic levels (words versus phrasal structures) was disrupted by noise, on the one hand, and language proficiency, on the other (low-frequency environmental noise during the recordings prevented us from analyzing entrainment to the sentential level). The MEG responses were transformed into the frequency domain, and we retained for subsequent analysis the 5 neural response components that were the most consistent over trials, as identified by spatial filters (see Methods). Results revealed the distinct influence of noise on the cortical tracking of different linguistic levels. A linear mixed effects regression on the amplitude of syllabic and phrasal peaks with Age of Acquisition, Language proficiency, Noise, and their interactions as continuous regressors revealed that while the level of tracking at the syllabic level was only affected by Noise (F(3,144) = 2.64; *p* = .05), at the phrasal level there was a reliable interaction between Noise and Language Proficiency (F(3,144) = 2.76; *p* = .04). This shows that the tracking capabilities of noisy signals varies across individuals depending on their proficiency. Further, there was a significant correlation between the amplitude of the syllabic peak, and that of the phrasal peak (*t*(202) = 40; *p* < .001), suggesting a relation between subjects’ capacity to segment incoming speech and parsing the syntactically-relevant structure.

Analogously to the behavioral data analysis, we complemented the continuous analysis with a categorical analysis where we assessed how individuals of different language profiles tracked both syllabic and phrasal structures. In consonance with the regression analysis, we found that participants of all proficiency combinations tracked the syllabic rhythm at all levels of noise, and the amplitude of this tracking response decreased as noise increased (Fig. 3; 3.2 Hz response in upper panels, 4 Hz response in lower panels). However, the tracking of phrasal structure was heavily dependent on language proficiency and comprehension (Fig. 3; 1.6 Hz response in upper panels, 2 Hz response lower panels).

**Fig. 3.**
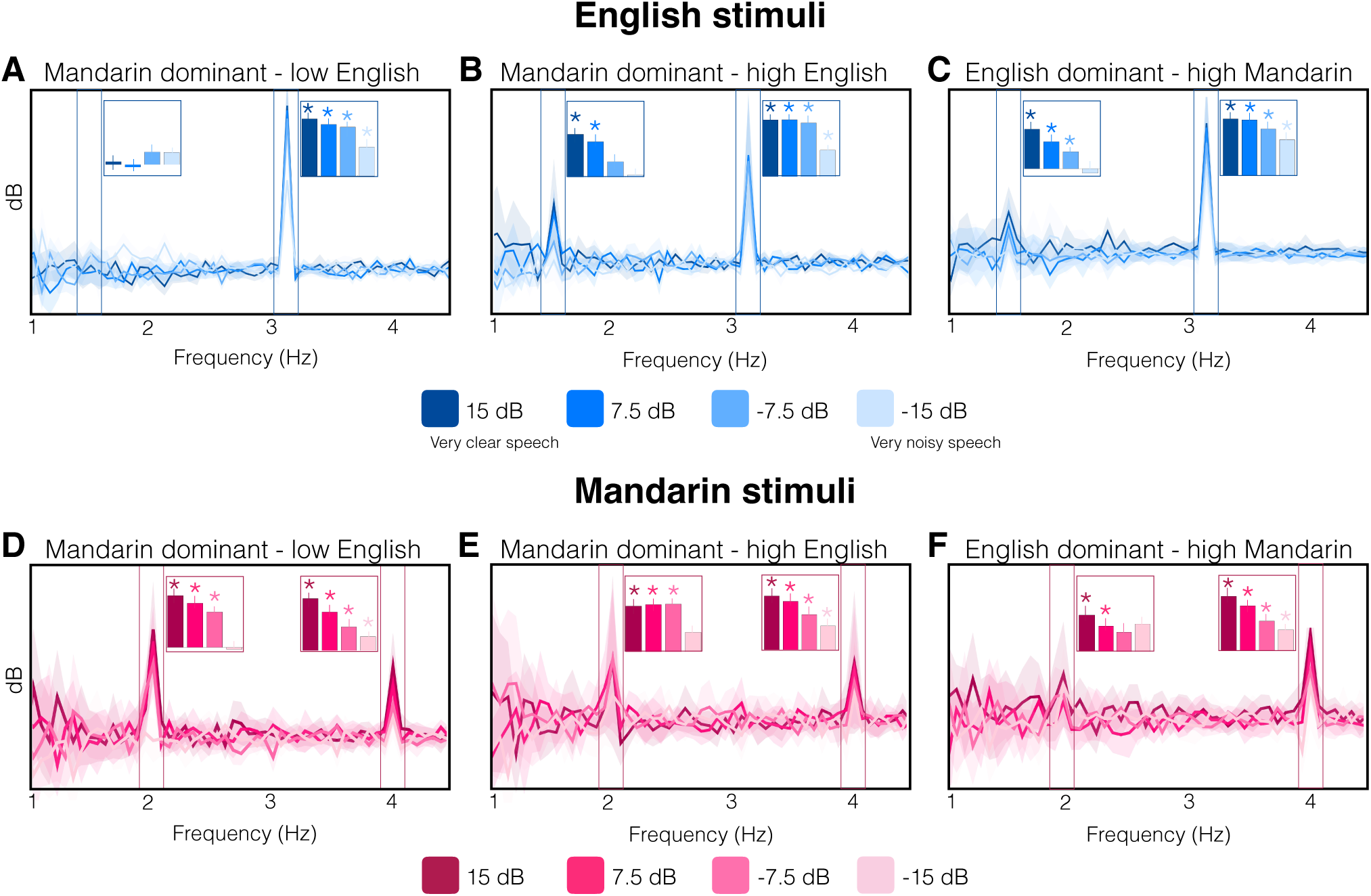
MEG-derived neural response spectra for each language group (Mandarin dominant – low English (n=16), Mandarin dominant – high English (n=12) and English dominant – high Mandarin (n=12). Solid lines indicate average response; shading is the 95% C.I. Spectral peaks at corresponding frequencies reflect whether there was neural tracking of syllabic or phrasal rhythms at a given level of noise. Frequency bins with significantly stronger power than two neighbors on each side are marked with a * of the corresponding color (p < 0.05, paired one-sided t-test, FDR corrected).

Specifically, with regard to speech rhythm tracking, Mandarin speakers with low English proficiency tracked syllabic rhythm in Mandarin (15 dB *p* < .001; 7.5 dB *p* = .002; −7.5 dB *p* = .001; −15 dB *p* = .008) and English (15 dB *p* < .001; 7.5 dB *p* < .001; −7.5 dB *p* = .003; −15 dB *p* = .002); as did Mandarin native speakers with high English proficiency (Mandarin: 15 dB *p* = .04; 7.5 dB *p* = .04; −7.5 dB *p* = .04; −15 dB *p* = .03; English: 15 dB *p* < .04; 7.5 dB *p* = .04; −7.5 dB *p* = .03; −15 dB *p* = .04) and English dominant speakers (English: all noise-levels *p* < .001; Mandarin: 15 dB and 7.5 dB *p* < .001; −7.5 dB *p* = .021; −15 dB *p* = .01).

In contrast, there was a clear disparity in the tracking response to phrase-level structure across languages depending on the proficiency combinations of the participants. Mandarin dominant speakers with low English proficiency did not track English phrases (15 dB *p* = .33; 7.5 dB *p* = .52; −7.5 dB *p* < .1 and −15 dB *p* = .21; Fig. 3A) although they did track phrases in Mandarin at all levels of noise (15 dB, 7.5 dB and −7.5 dB *p* < .001) except during pure noise (−15 dB *p* = .53; Fig. 3D). As proficiency in English increased, so did the tracking of the English phrases. Mandarin speakers with high English proficiency did track phrases at the clearest levels of speech in English (15 dB *p* < .04; 7.5 dB *p* = .02) although not at the two noisier levels (−7.5 dB *p* = .16 and −15 dB *p* = .22; Fig. 3B); while also being able to track Mandarin phrases at all levels of noise (15 dB *p* = .04; 7.5 dB *p* = .04; and −7.5 dB *p* = .02) except during pure noise (−15 dB *p* = .15; Fig. 3E). Finally, English dominant speakers showed phrasal tracking of English phrases at 15 and 7.5 dB, as did the Mandarin dominant speakers with high English proficiency (*p* = .04 and *p* = .002, respectively). But crucially, the speakers with higher English proficiency were additionally able to track phrases at −7.5 dB (*p* = .04; Fig. 3C). Hence the increase in English proficiency was accompanied by tracking of phrasal structures at higher levels of noise. In contrast, these same English dominant speakers whose Mandarin was mildly worse were only able to track phrases in Mandarin at the two clearest levels of speech (15 dB (*p* = .03) and 7.5 dB (*p* = .01)) but not at −7.5 dB (*p* = .09) or −15 dB (*p* = .1) like Mandarin dominant speakers had done (Fig. 3F). Hence, the categorical analyses confirmed what the regression analysis revealed: noise and proficiency interact critically in the tracking of higher-level linguistic structures.

These results build on previous findings reporting an effect of intelligibility on cortical tracking of speech (Doelling et al., 2014; Park, Ince, Schyns, Thut & Gross, 2015; Peelle, Gross, & Davis, 2012), but reveal a more complex and informative pattern than previously known. Specifically, we show (i) that not all levels of entrainment are affected equally by noise and (ii) that not only the physical properties of the stimuli but also the language proficiency of the listener affects the degree of entrainment to different linguistic structures. These results complement research showing that overall neural oscillatory activity underlying speech processing also varies with second language proficiency (Pérez, & Duñabeitia, 2019; Pérez, Carreiras, Dowens, & Duñabeitia, 2015).

### MEG: Source localization

Finally, we performed analyses to identify from where in the cortex the reported activation patterns were emerging. We source-localized the same 5 MEG components submitted to the frequency-domain analysis and compared neural responses to English and Mandarin at −7.5 dB, as this was the SNR at which language proficiency clearly determined the presence or absence of phrasal entrainment. The analysis revealed that activity in areas surrounding the auditory cortex elicited increased activity in response to the better-understood language. Furthermore, this increase widened commensurate with the imbalance between languages (Fig. 4). This is consistent with previous research that localized intelligibility effects (Peelle, Gross, & Davis, 2012; Keitel, Gross, & Kayser, 2018).

**Fig. 4.**
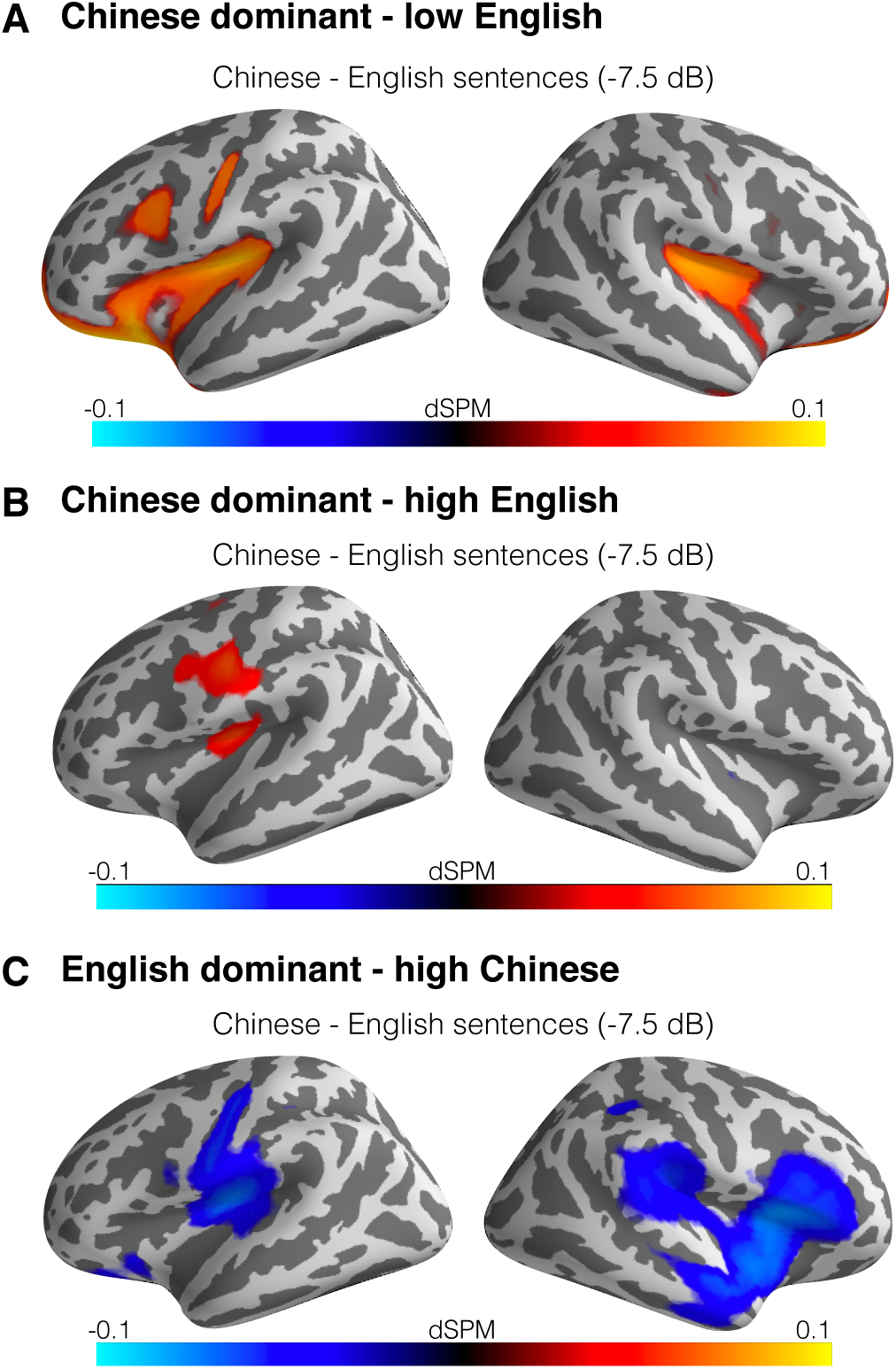
Whole brain source localization of the 5 neural response components most consistent over trials, identified with a Denoising Source Separation (DSS) technique. The whole brain images show the result of subtracting the average activity elicited by English sentences (at −7.5 dB) from the average activity elicited by Mandarin sentences (at −7.5 dB), averaged across subjects. Activity is displayed in noise-normalized statistical parametric maps (dSPM; blue indicates higher activity for English stimuli, red for Mandarin stimuli).

## Discussion

The combined behavioral and neurophysiological data we present capitalize on recent findings and illuminate speech and language processing in new ways. In particular, it has been shown that neural entrainment to speech signals enables both tracking of the acoustic input, principally the amplitude envelope, but also higher order structure building operations that are not accessible from the physical input alone (Fig.1, Fig.3; Ding et al., 2016; Luo & Poeppel, 2007, Keitel et al., 2018). In this study, we found that neural responses at the theta band that track the physical speech rhythm are only affected by noise – but not by language proficiency. In contrast, neural tracking of phrasal structure at delta level was affected by the interaction of noise and language knowledge. We advance current understanding in two significant ways, deriving from the parametric nature of the experimental design, in which we concurrently vary the quality of the speech signal (Fig.1) and the language proficiency of the listeners (Fig.2).

First, the paradigm allows us to establish that the language knowledge of the listener determines the spectral information required for speech recognition (cf. Shannon, Zeng, Kamath, Wygonski, & Ekelid, 1995). In other words: there is no such thing as a categorical limit on how impoverished the signal can be before comprehension is compromised. Instead, this boundary is malleable and shifts in concordance with the linguistic capabilities of the listener.

Relatedly, we show that neurophysiologically it is at the phrasal level (cf. Fig. 1) that differential knowledge of language is especially influential. There is a minimum level of SNR (−7.5 dB in native language and 7.5 dB in the non-native language; Fig. 3) necessary to facilitate phrase-level tracking, which in turn forms the basis for structure building. Listeners who track their L1 well at −7.5 dB fail at tracking their L2 at that same SNR, suggesting that the tracking impairment is not due to peripheral causes but implicates higher-order limitations. This finding suggests that it is not only conscious comprehension but also unconscious neural processes that are sensitive to the interaction between noise and language knowledge. Importantly, this result is consistent with recent findings showing that delta band tracking yields a significant prediction of speech comprehension (Etard & Reichenbach, 2019), and with research suggesting that delta oscillations are not primarily involved in early sound analysis and phonological processing, but rather they reflect the encoding of abstract syntactic structures (Kosem & van Wassenhove, 2016).

These results shed new light on the conceptualization of multilingual language comprehension. While previous accounts revealed that the source of the prevalent bilingual impairment to comprehend speech in noise emerges from deficiencies in access to lexical or syntactic information in this language (Ferreira et al., 2009; Hahne & Friederici, 2001; Hervais-Adelman et al., 2014), our results suggest that this impairment is additionally reflected, and perhaps instigated by, a failure to successfully complete a lower-level processes. Although, this experiment cannot prove causality by itself, this proposal is supported by recent research in monolingual individuals showing that the capability to entrain to linguistic structures can in fact causally affect comprehension (Zoefel et al., 2018).

Mechanistically, by hypothesis, comprehension may be enhanced by a feedback loop such that bottom-up rhythmic structure and top-down information mutually aid the prediction and processing of upcoming signals (Peelle et al., 2012). In bilinguals, both of these processes may be compromised: While previous research has focused on the information availability aspect (Hervais-Adelman et al., 2014; Golestani, Hervais-Adelman, Obleser, & Scott, 2013), we show that the impoverished comprehension of speech in noise by L2 learners is *also* determined by a disruption in the entrainment to linguistic structures. Importantly, these processes do not seem dissociable: there was a significant correlation between the amplitude of the syllabic peak and the amplitude of the phrasal peak, suggesting a meaningful relation between subjects’ capacity to segment incoming speech (i.e., a signal-based low-level process) and parsing the syntactically-relevant structure (i.e., a high-level process).

In sum, our results characterize the minimum SNR requirements for neural entrainment to different linguistic structures, specify the differential influence of noise and knowledge on syllabic and phrasal tracking, and reveal a neurophysiological pattern that may underlie the widely experienced phenomenon of compromised comprehension of second language speech in noisy environments.

## Acknowledgments: Funding

National Institute of Health grant 5R01DC05660 and Max Planck Society funding to D.P, NYUAD Institute grant G1001 to L.P., Woodcock Institute Research Award to E.B.E.

## Author contributions

E.B.E and D.P designed the experiment; E.B.E collected and performed data analyses, N.D. supported data analyses; E.B.E, D.P and L.P wrote the manuscript.

## Competing interests

Authors declare no competing interests.

## Data and materials availability

All data and analysis code is available upon request.

---

**Supplementary Figure 1:**
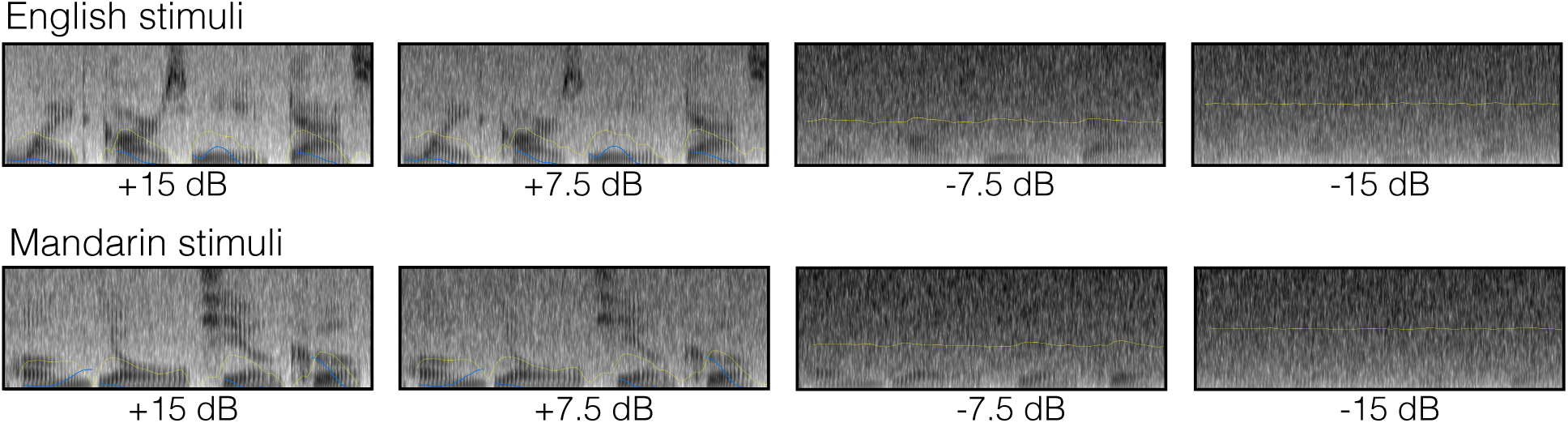
Spectrograms for a sample stimuli sentence at each SNR. Top row: spectrograms for English stimuli. Bottom row: spectrograms for Mandarin stimuli.

